# Myogenetic oligodeoxynucleotide (myoDN) complexed with berberine promotes differentiation of chicken skeletal muscle myoblasts

**DOI:** 10.1101/2020.12.19.423622

**Authors:** Yuma Nihashi, Sayaka Shinji, Koji Umezawa, Takeshi Shimosato, Tamao Ono, Hiroshi Kagami, Tomohide Takaya

## Abstract

Skeletal muscle myoblasts are myogenic precursors that develop into myofibers during muscle formation. Improvement of myoblast differentiation is important for advancing meat production by domestic animals. We recently identified novel oligodeoxynucleotides (ODNs) termed myogenetic ODNs (myoDNs) that promote the differentiation of mammalian myoblasts. An isoquinoline alkaloid, berberine, forms a complex with one of the myoDNs, iSN04, and enhances its activities. This study investigated the effects of myoDNs on chicken myoblasts to elucidate their species-specific actions. Seven myoDNs (iSN01-iSN07) were found to facilitate the differentiation of chicken myoblasts into myosin heavy chain (MHC)-positive myocytes. The iSN04-berberine complex exhibited a higher myogenetic activity than iSN04 alone, which was shown to enhance the differentiation of myoblasts into myocytes, myotube formation, and myogenic gene expression (MyoD, myogenin, MHC, and myomaker). These data indicate that myoDNs promoting chicken myoblast differentiation may be used as potential feed additives in broiler diets.

## 1. Introduction

The demand for chicken meat has increased worldwide over the past half-century [1]. To improve meat production, breeding and feedstuff of chickens have been intensively advanced. Genetic selection has established broiler chickens whose muscle growth rates are several times higher than those of other breeds such as layer chickens [2]. Skeletal muscle tissue is composed of numerous myofibers that are multi-nucleated giant myocytes. Each myofiber has dozens of stem cells, known as satellite cells. During muscle growth, satellite cells are activated to the proliferative myogenic progenitor cells, termed myoblasts. After several rounds of cell division, myoblasts differentiate into contractile myocytes. Eventually, myocytes fuse to existing myofibers or fuse with each other to form myotubes, which subsequently outgrow to nascent myofibers [3]. Therefore, the characteristics of myoblasts reflect the muscle phenotype of animals. We previously reported that broiler myoblasts actively proliferate, promptly differentiate into myotubes, and present differential gene expression patterns compared to layer myoblasts [4, 5]. This corresponds well with broiler phenotype and suggests that enhancement of myogenic differentiation of broiler myoblasts accelerates muscle development and shortens the rearing period.

To maximize the performance of broilers, various diets and feed additives have been developed. Synthetic amino acids, processed plant proteins, and dried plasma have been utilized as sources for chicken nutrition [6]. Accurate calcium concentration in broiler diets is essential for increasing feed intake and body weight [7]. Supplementation of transgenic phytase improves body weight and upregulates genes involved in growth response and meat quality in leg muscles [8]. However, there is no diet or additive that directly acts on chicken myoblasts.

We recently identified a series of 18-base oligodeoxynucleotides (ODNs) termed myogenetic ODNs (myoDNs) that induce extensive differentiation of murine and human myoblasts independently of Toll-like receptors (TLRs) [9]. One of the myoDNs, iSN04, is spontaneously incorporated into myoblasts, antagonizes nucleolin to increase p53 protein levels, and modulates gene expression to lead to myogenic fate. Intriguingly, iSN04 forms a complex with an isoquinoline alkaloid, berberine. The iSN04-berberine complex exhibits higher myogenetic activity than iSN04 alone. This is probably because berberine shifts the iSN04 conformation to a stable and active form [9]. myoDNs are the first instance of ODNs promoting myoblast differentiation, which may contribute to improving meat production by domestic animals. To apply myoDNs to a broad variety of animals in the future, the effects of myoDNs on non-mammalian myoblasts should be validated.

Assessment of species-specificity is technically and industrially important for ODN application. In the field of TLR-dependent immunogenic ODNs, for example, ODN-2006 stimulates both murine and human TLR9, but ODN-1826 is recognized only by murine TLR9 [10]. Since aves express TLR21 instead of TLR9, a novel type of immunogenic ODNs was identified using chicken macrophages [11]. These studies suggest that myoDNs may possess species-specificy. Amino acid sequences of the iSN04-target protein, nucleolin, are relatively but not extremely similar between aves and mammals (62.5% identity between chicken and human, 59.6% identity between chicken and mouse). This study investigated the myogenetic effects of myoDNs on broiler myoblasts to reveal their secies-specific actions and potential availabilities on chicken myoblasts.

## 2. Materials and methods

### 2.1. Chemicals

Phosphorothioated (PS)-ODNs (Fig. 2B) were synthesized and purified via HPLC (GeneDesign, Osaka, Japan). PS-ODNs and berberine hydrochloride (Nacalai, Osaka, Japan) were dissolved in endotoxin-free water. An equal volume of endotoxin-free water containing no PS-ODNs or berberine served as a negative control. For iSN04-berberine complex formation, iSN04 and berberine were pre-mixed in RPMI1640 medium (Nacalai) and incubated at 20°C for 30 min.

**Fig 1.**
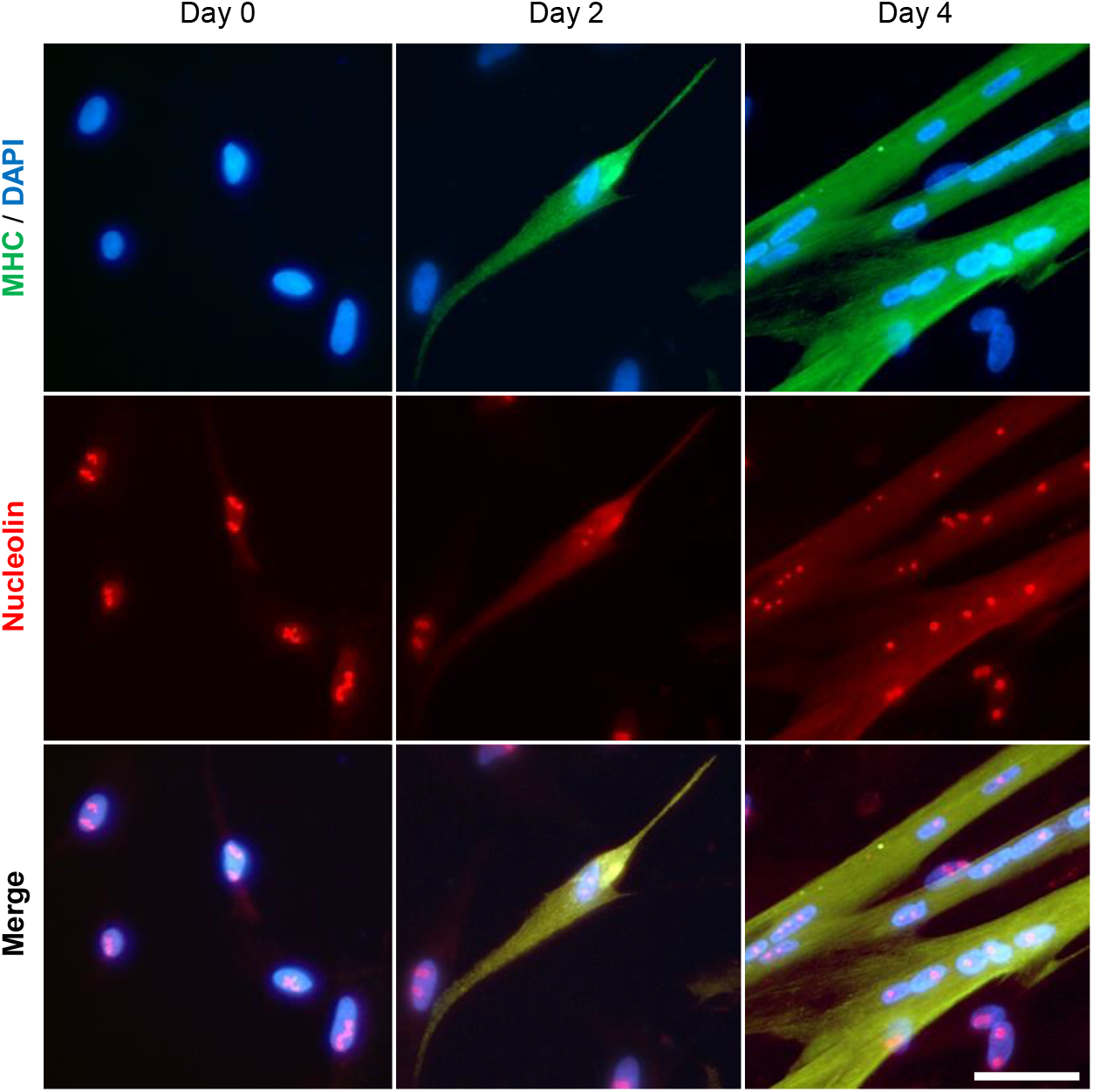
Nucleolin localization in chicken myoblasts during differentiation. Representative images of nucleolin and MHC staining of the myoblasts cultured in DM at days 0, 2, and 4. Scale bar, 50 μm.

**Fig 2.**
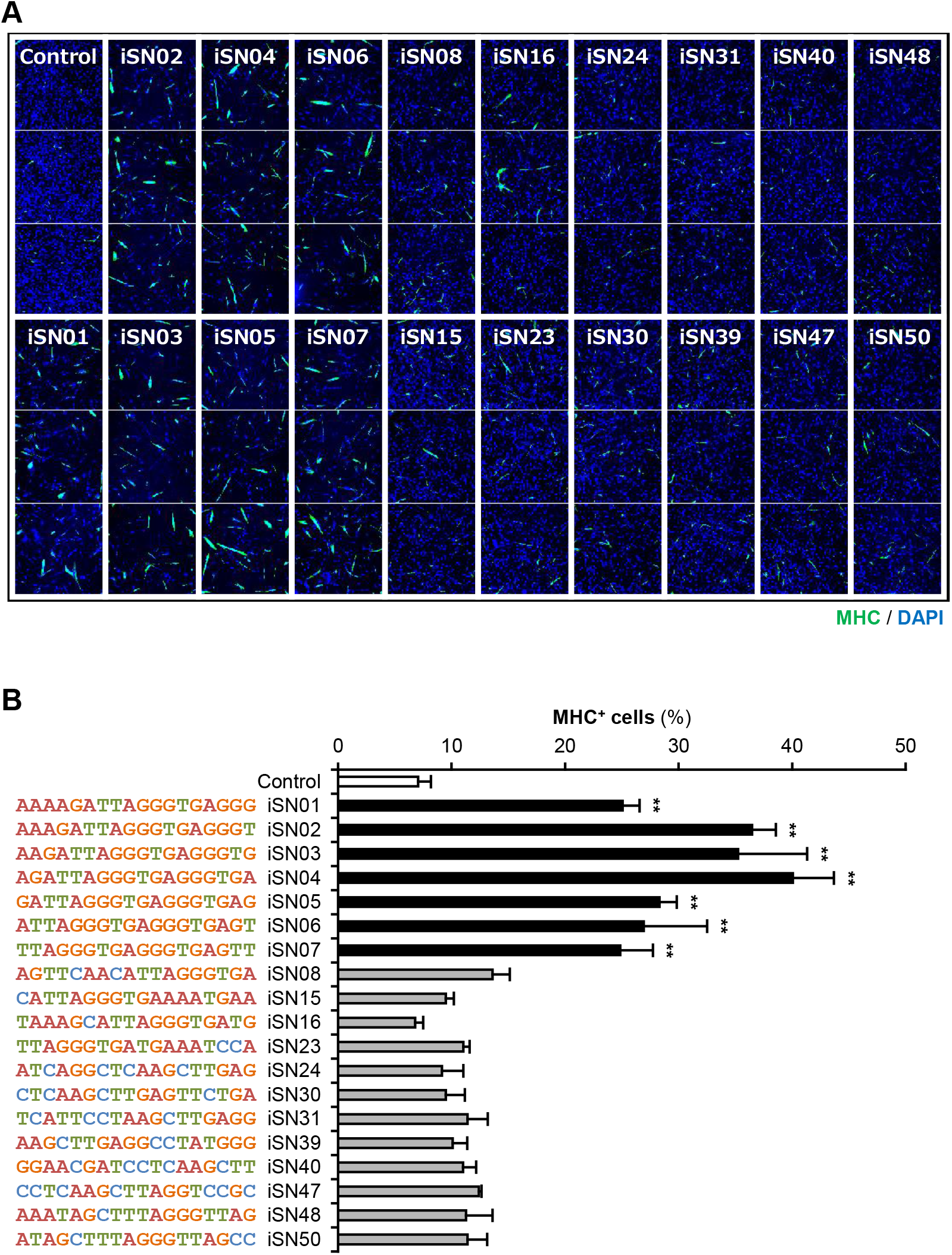
Screening of PS-ODNs on the differentiation of chicken myoblasts. (A and B) Immunofluorescent images of MHC and DAPI staining (A) and the ratio of MHC^+^ cells (B). Chicken myoblasts were treated with 10 μM PS-ODN in GM for 48 h. ** *p* < 0.01 vs control (Dunnett’s test). *n* = 3.

### 2.2. Isolation and culture of chicken myoblasts

All experimental procedures were conducted in accordance with the Regulations for Animal Experimentation of Shinshu University, and the animal protocol was approved by the Committee for Animal Experiments of Shinshu University. Chicken skeletal muscle myoblasts were isolated from the leg muscles of E10 embryos as previously described in detail [4, 12, 13]. The myoblasts of Barred Plymouth Rock chicken (Goto Poultry Farm, Gifu, Japan) were used for nucleolin staining and PS-ODN screening, and those of UK Chunky chicken (National Federation of Agricultural Cooperative Associations, Tokyo, Japan) were used for subsequent experiments. The myoblasts were cultured at 37°C under 5% CO2 on the dishes or plates coated with collagen type I-C (Cellmatrix; Nitta Gelatin, Osaka, Japan) throughout the experiments. The myoblasts were maintained in a growth medium (GM) consisting of RPMI1640, 20% fetal bovine serum (FBS) (HyClone; GE Healthcare, UT, USA), 1% non-essential amino acids (Wako, Osaka, Japan), 1% chicken embryo extract (US Biological, MA, USA), 2 ng/ml basic fibroblast growth factor (Wako), and a mixture of 100 units/ml penicillin and 100 μg/ml streptomycin (PS) (Nacalai). To induce myogenic differentiation, GM was replaced with a differentiation medium (DM) consisting of DMEM (Nacalai), 2% FBS, and PS after 24 h of seeding myoblasts.

### 2.3. Immunocytochemistry

Myoblasts (5.0×10^3^ cells/well) were seeded on 96-well plates for screening, or 1.0×10^5^ myoblasts/dish were seeded on 30-mm dishes for high-resolution imaging. The next day, the medium was replaced with GM containing 10 μM PS-ODN and 10 μM berberine. Immunostaining was performed as previously described [4, 12, 13]. The myoblasts were fixed with 2% paraformaldehyde, permeabilized with 0.2% Triton X-100 (Nacalai), and immunostained with 0.5 μg/ml mouse monoclonal anti-myosin heavy chain (MHC) antibody (MF20; R&D Systems, MN, USA) and 1.0 μg/ml rabbit polyclonal anti-nucleolin antibody (ab22758; Abcam, Cambridge, UK). Cell nuclei were stained with DAPI (Nacalai). Fluorescent images for screening were automatically captured using CellInsight NXT (Thermo Fisher Scientific, MA, USA). The ratio of MHC^+^ cells was calculated using HCS Studio: Cellomics Scan software (Thermo Fisher Scientific). High-resolution fluorescent images were taken under an EVOS FL Auto microscope (AMAFD1000; Thermo Fisher Scientific). The ratio of MHC^+^ cells was defined as the number of nuclei in all MHC^+^ cells divided by the total number of nuclei, and the fusion index was defined as the number of nuclei in multinuclear MHC^+^ myotubes divided by the total number of nuclei using ImageJ software (National Institutes of Health, USA).

### 2.4. Cell counting

Myoblasts (2.5×10^4^ cells/well) were seeded on 12-well plates. After 24 h, the myoblasts were treated with 3 or 10 μM iSN04. For counting, the myoblasts were completely dissociated by treatment with 0.25% trypsin with 1 mM EDTA (Wako) at 37°C for 5 min. The number of myoblasts was counted using a hemocytometer. The dissociated cells were not seeded again.

### 2.5. EdU staining

EdU (5-ethynyl-2’-deoxyuridine) staining was performed as previously described [4]. Myoblasts (1.0×10^5^ cells/dish) were seeded on 30-mm dishes. The next day, the myoblasts were treated with 0 or 3 μM iSN04 for 48 h and then treated with 10 μM EdU for 3 h. EdU was stained using Click-iT EdU Imaging Kit (Thermo Fisher Scientific). Cell nuclei were stained with DAPI. The ratio of EdU^+^ cells was defined as the number of EdU^+^ nuclei divided by the total number of nuclei using ImageJ software.

### 2.6. Quantitative real-time RT-PCR (qPCR)

Myoblasts (3.0×10^5^ cells/dish) were seeded on 60-mm dishes. After 24 h, the myoblasts were treated with 10 μM of iSN04 and berberine for 8 or 24 h. Total RNA from the myoblasts was isolated using NucleoSpin RNA Plus (Macherey-Nagel, Düren, Germany). RNA was reverse transcribed using ReverTra Ace qPCR RT Master Mix (TOYOBO, Osaka, Japan). qPCR was performed using GoTaq qPCR Master Mix (Promega, WI, USA) with StepOne Real-Time PCR System (Thermo Fisher Scientific). The amount of each transcript was normalized to that of tyrosine 3-monooxygenase/tryptophan 5-monooxygenase activation protein zeta (*YWHAZ*). The results are presented as fold-change. Primer sequences are listed in Table 1.

**Table 1.**
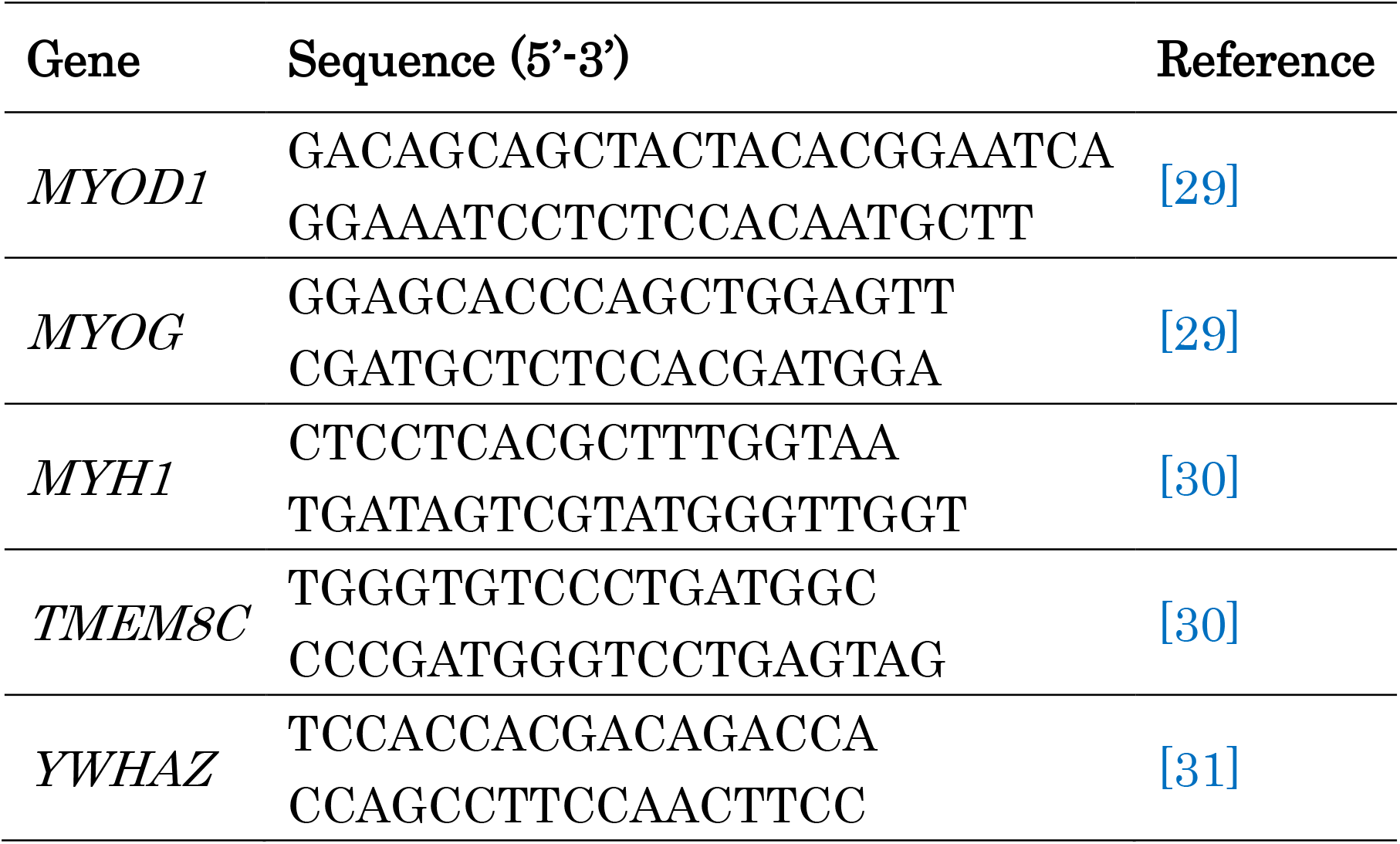
Primer sequences for qPCR.

### 2.7. Statistical analyses

Results are presented as mean ± standard error. Statistical comparisons were performed using multiple comparison test with Dunnett’s test, Williams’ test, or Scheffe’s *F* test, where appropriate following one-way analysis of variance. Statistical significance was set at *p* < 0.05.

## 3. Results

### 3.1. myoDNs promote myogenic differentiation of chicken myoblasts

Although nucleolin is a target protein of myoDNs [9], its expression in chicken myoblasts has not been reported. Immunostaining of primary-cultured chicken myoblasts indicated that nucleolin was localized in the nuclei and was especially concentrated in the nucleoli of undifferentiated myoblasts (Fig. 1, day 0). In the differentiated MHC^+^ mononuclear myocytes and multinuclear myotubes, nucleolin remained in the nucleoli and diffused into the cytoplasm (Fig. 1, days 2 and 4). The shift of nucleolin localization during chicken myoblast differentiation corresponded well to that observed in mice and humans [9]. Next, nineteen 18-base PS-ODNs, including seven myoDNs (iSN01-iSN07), were screened for their effects on myogenic differentiation of chicken myoblasts. The myoblasts maintained in GM were treated with 10 μM PS-ODNs for 48 h and immunostained for MHC (Fig. 2A). iSN01-iSN07 significantly increased the ratio of MHC^+^ cells, but other PS-ODNs did not affect the differentiation of myoblasts (Fig. 2B). This indicates that myoDNs can promote myogenic differentiation of chicken myoblasts, as it does in murine and human myoblasts. Since iSN04 exhibits the highest myogenetic activity in chicken myoblasts as well as in murine myoblasts [9], iSN04 was utilized in the following experiments.

### 3.2. iSN04 suppresses proliferation of chicken myoblasts

Proliferation and differentiation are inverse processes and negatively regulate each other in stem cells and precursor cells [14]. In murine myoblasts, iSN04 represses cell proliferation by promoting myogenic differentiation [9]. Continuous cell counting revealed that iSN04 also suppressed the growth of chicken myoblasts in a dose-dependent manner. The number of chicken myoblasts treated with 3 or 10 μM iSN04 for 48 h was significantly lower than that of the control group (Fig. 3A). DNA replication in myoblasts was measured by EdU staining (Fig. 3B). The ratio of EdU^+^ cells was significantly decreased in chicken myoblasts treated with 10 μM iSN04 for 48 h (Fig. 3C). These data indicate that iSN04 suppresses the proliferation of chicken myoblasts.

**Fig 3.**
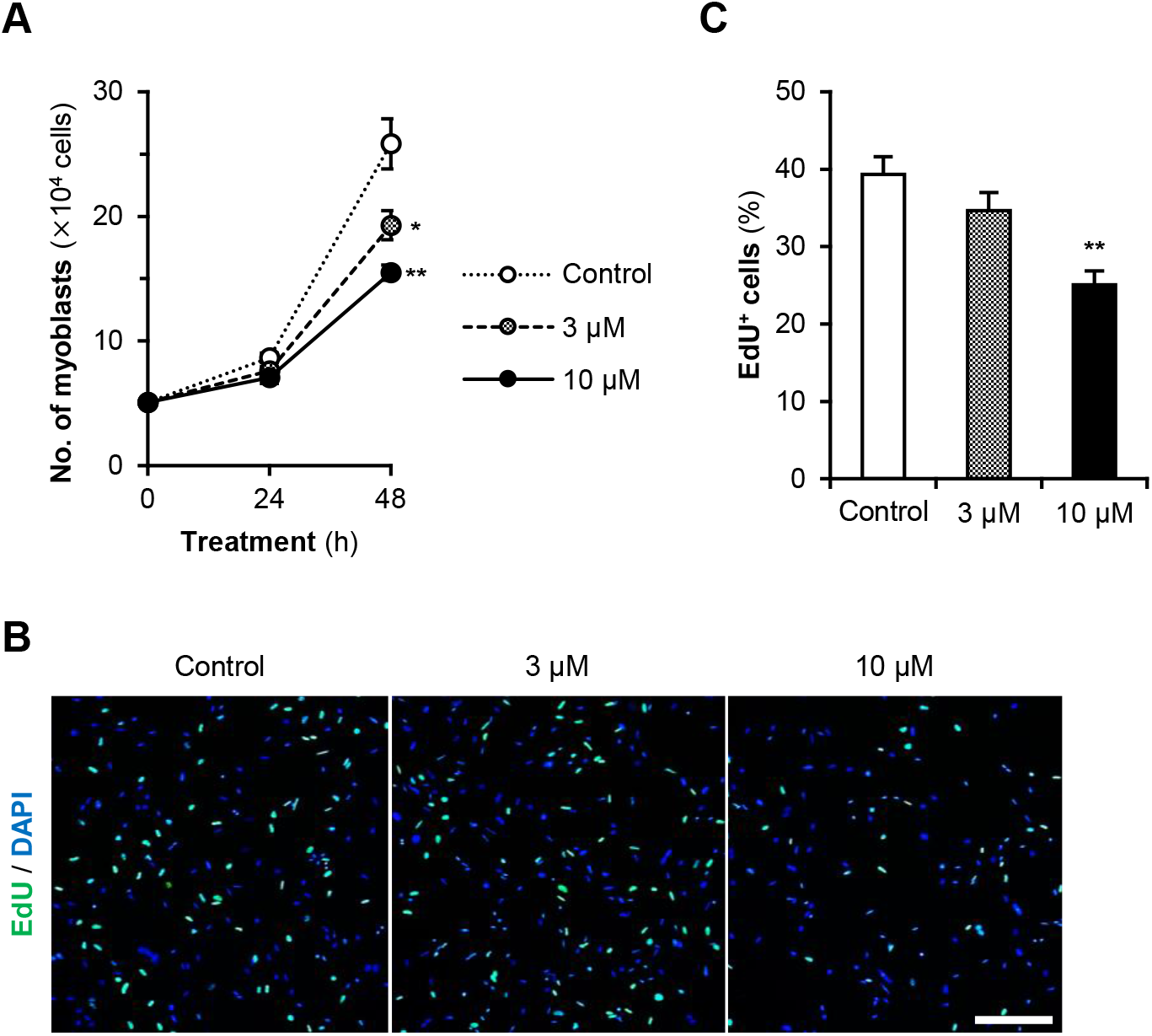
Effects of iSN04 on the growth of chicken myoblasts. (A) The numbers of the myoblasts treated with 3 or 10 μM iSN04 in GM. * *p* < 0.05, ** *p* < 0.01 vs control at 48 h (Williams’ test). *n* = 3. (B and C) Representative images of EdU and DAPI staining (B) and the ratio of EdU^+^ myoblasts (C). Chicken myoblasts were treated with 3 or 10 μM iSN04 in GM for 48 h. Scale bar, 200 μm. ** *p* < 0.05 vs control (Williams’ test). *n* = 4.

### 3.3. Berberine enhances myogenetic activity of iSN04 on chicken myoblasts

An isoquinoline alkaloid, berberine, physically interacts with iSN04 via the 13-15th guanines and forms a tight complex. In murine myoblasts, the iSN04-berberine complex exhibits a higher myogenetic activity than iSN04 alone, probably because of stabilized and optimized conformation [9]. To examine whether berberine enhances iSN04 activity in chicken, 10 μM of berberine, iSN04, or the pre-mixed iSN04-berberine complex were administered to chicken myoblasts for 48 h (Fig. 4A). Consistent with the screening results, iSN04 significantly increased the ratio of MHC^+^ cells (23.9%) and fusion index (18.7%) compared to those of the control (8.0% and 5.2%, respectively) (Fig. 4B&C). The iSN04-complex significantly promoted the differentiation of myoblasts into MHC^+^ cells (42.4%) and multinuclear myotubes (32.0%), and its effects were significantly higher than that of iSN04 alone. As berberine alone did not alter the differentiation of chicken myoblasts, it was confirmed that the enhanced activity of the iSN04-berberine complex is not a synergistic effect. These data indicate that berberine enhances the myogenetic activity of iSN04 in chicken myoblasts, similar to that in murine myoblasts.

**Fig 4.**
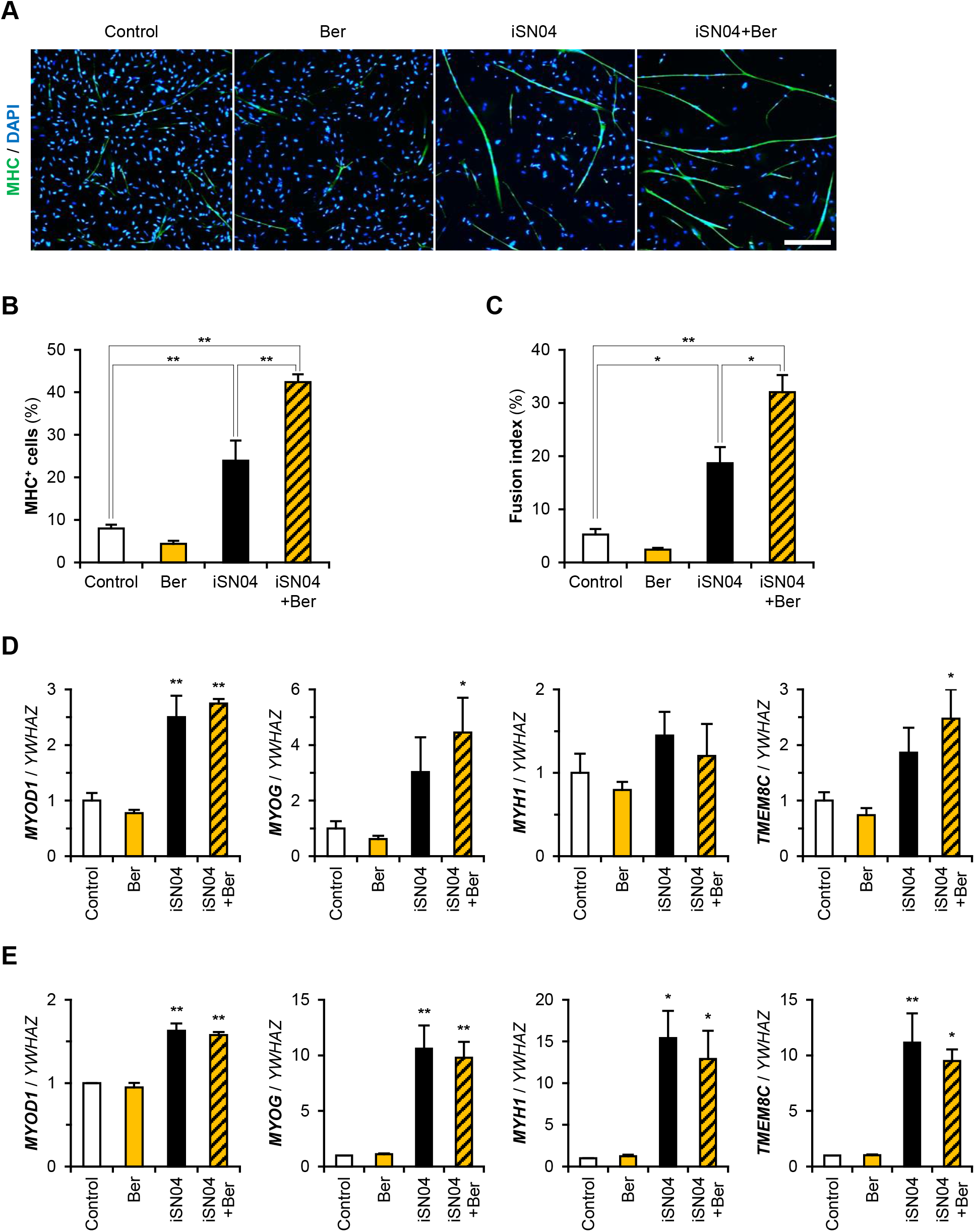
Effects of the iSN04-berberine complex on the differentiation of chicken myoblasts. (A-C) Representative images of MHC and DAPI staining (A), the ratio of MHC^+^ cells (B), and fusion index (C). Chicken myoblasts were treated with 10 μM of berberine (Ber) and iSN04 in GM for 48 h. Scale bar, 200 μm. * *p* < 0.05, ** *p* < 0.01 (Scheffe’s *F* test). *n* = 4. (D and E) qPCR results of myogenic gene transcriptions in the chicken myoblasts treated with 10 μM of iSN04 and berberine in GM for 8 h (D) and 24 h (E). The mean value of the control was set to 1.0 for each gene. * *p* < 0.05, ** *p* < 0.01 vs control (Dunnett’s test). *n* = 3-4.

### 3.4. iSN04 upregulates myogenic gene expression in chicken myoblasts

To reveal the effects of iSN04 and the iSN04-berberine complex on gene expression in chicken myoblasts, transcription levels of MyoD (*MYOD1*), myogenin (*MYOG*), skeletal muscle MHC (*MYH1*), and myomaker (*TMEM8C*; the muscle-specific membrane protein required for myotube formation) were quantified by qPCR. At 8 h after the administration, iSN04 and the iSN04-complex significantly upregulated MyoD expression, which is the initial step of myogenic differentiation (Fig. 4D). Furthermore, the iSN04-complex, but not iSN04 alone, significantly induced myogenin and myomaker transcription. This indicates the higher myogenetic activity of the iSN04-complex rather than that of iSN04 alone. At 24 h after the treatment, both iSN04 and the iSN04-berberine complex enhanced the transcription of MyoD, myogenin, MHC, and myomaker to the same extent (Fig. 4E). In both timepoints, berberine did not alter any gene expression. These data demonstrate that iSN04, especially the iSN04-berberine complex, starts to modulate gene expression programs in chicken myoblasts shortly after the administration.

## 4. Discussion

The present study showed that a series of 18-base telomeric ODNs termed myoDNs (iSN01-iSN07) promote the myogenic differentiation of chicken myoblasts. Among the myoDNs, iSN04 exhibited the highest activity in chicken myoblasts. These data corresponds well with our previous results using murine and human myoblasts [9], ensuring that myoDNs are available to induce differentiation of both avian and mammalian myoblasts. Myoblast differentiation and myotube formation are essential processes during the development, formation, and regeneration of skeletal muscle tissue [3]. Thus, the myoblast properties are closely related to the muscle phenotype of the animals. Broiler chicken myoblasts show potent differentiation ability compared to layer chicken myoblasts [4]. However, the inherent differentiation ability of broiler myoblasts diminishes with age and body weight [15]. The pectoralis major muscle of adult broilers frequently displays myopathic wooden breast syndrome characterized by degenerative myofibers, fibrosis, and mitotic myoblasts [16, 17]. Therefore, accelerating myoblast differentiation is anticipated to facilitate muscle development and eventually improve the meat quality of chickens. myoDNs are single-strand short ODNs that directly target myoblasts. ODNs are chemically synthesized, stable, and safe molecules that have been utilized as nucleic acid drugs. These features of myoDNs make them suitable for use as beneficial dietary additives for domestic fowls.

The iSN04-berberine complex further promoted myogenic differentiation and myotube formation in chicken myoblasts compared to iSN04 alone. Berberine is a safe isoquinoline alkaloid derived from medicinal plants and exhibits multiple bioactivities such as anti-inflammatory and anti-tumor effects [18, 19]. It has been utilized in several clinical trials [20]. In addition, berberine is a ligand of telomeric DNA, and stabilizes its conformation [21]. We previously reported that berberine physically interacts with iSN04 via the 13-15th guanines and modulates iSN04 conformation. As a result, the iSN04-complex exerts a potent myogenetic effect on murine myoblasts [9]. The present study confirmed that the berberine-enhanced iSN04 activity can also be observed in aves. The potentiation of myoDNs using berberine is a valuable technology to refine ODN functions.

iSN04 physically interacts with a multifunctional protein, nucleolin [9]. One of the functions of nucleolin is binding to p53 mRNA to interfere with translation [22, 23]. It has been widely known that the p53 signaling pathway promotes myogenic differentiation [24-26]. In human myoblasts, iSN04 antagonizes nucleolin and increases p53 protein levels. Then, the iSN04-activated p53 signaling pathway induces myogenic differentiation with coupled cell cycle arrest by upregulating the expression of muscle genes including MyoD and myogenin [9]. In the present study, iSN04 promoted differentiation and suppressed proliferation of chicken myoblasts, strongly suggesting that the action mechanism of iSN04 is common between avian and mammalian myoblasts. The actions of ODNs are often species-specific. Unmethylated CpG-ODNs are ligands for TLR9, but their optimal motifs are different between mice and humans [27]. The specificity of CpG-ODNs is probably dependent on binding affinities to the ectodomain of TLR9 in each species [10]. In contrast, iSN04 showed identical effects among chicken, murine, and human myoblasts. iSN04 is considered to interact with four RNA-binding domains (RBD1-RBD4) of nucleolin because the established anti-nucleolin DNA aptamer AS1411 binds to these RBDs [28]. We have already confirmed that AS1411 also promotes mammalian myoblasts as well as iSN04 [9]. The amino acid sequences of chicken RBDs are highly homologous to those of mice and humans (RBD1 is 55.8% and 55.8%, RBD2 is 64.0% and 68.0%, RBD3 is 66.7% and 70.7%, and RBD4 is 89.5% and 88.2% identical to mice and humans, respectively). Sequence conservation, especially in RBD4 of nucleolin, is one of the reasons why iSN04 exerts myogenetic action on both avian and mammalian myoblasts.

In conclusion, myoDNs, including iSN04, promoted differentiation of chicken myoblasts to a similar extent as mammalian myoblasts. The myogenetic activity of iSN04 on chicken myoblasts was further enhanced by forming a complex with berberine. myoDNs directly targeting chicken myoblasts will be useful as dietary additives to improve meat production.

## Author contributions

TT designed the study. TT and YN wrote the manuscript. YN, SS, and TT performed experiments. TS designed and provided the ODNs. HK provided chicken embryos. KU and TO supervised the study.

## Acknowledgments

We thank Ms. Chikako Miyazaki for her excellent technical assistance. This study was supported in part by Grants-in-Aid from the Japan Society for the Promotion of Science to TT (19K05948) and HK (18K05939); Grants-in-Aid from the Kieikai Research Foundation (2018C002), the Ito Foundation, and the Shinshu Foundation for Promotion of Agricultural and Forest Science to TT; a Research Fellowship for Young Scientists from the Japan Society for the Promotion of Science (19J20888) and a Grant-in-Aid from the Fund of Nagano Prefecture to Promote Scientific Activity to YN (H29-3-12).

## References

[1] J.A. Arthur, G.A.A. Albers, Industrial perspective on problems and issues associated with poultry breeding, in W.M. Muir, S.E. Aggrey (Eds.), Poultry Genetics, Breeding and Biotechnology, CAB International, 2003, pp. 1–12. https://doi.org/10.1079/9780851996608.0001.

[2] G.N. Scheuermann, S.F. Bilgili, S. Tuzun, D. R. Mulvaney, Comparison of chicken genotypes: myofiber number in pectoralis muscle and myostatin ontogeny, Poult. Sci. 83 (2004) 1404–1412. https://doi.org/10.1093/ps/83.8.1404.

[3] N.A. Dumont, C.F. Bentzinger, M.C. Sincennes, M.A. Rudnicki, Satellite cells and skeletal muscle regeneration, Comp. Physiol. 5 (2015) 1027–1059. https://doi.org/10.1002/cphy.c140068.

[4] Y. Nihashi, K. Umezawa, S. Shinji, Y. Hamaguchi, H. Kobayashi, T. Kono, T. Ono, H. Kagami, T. Takaya, Distinct cell proliferation, myogenic differentiation, and gene expression in skeletal muscle myoblasts of layer and broiler chickens,Sci. Rep. 9 (2019) 16527. https://doi.org/10.1038/s41598-019-52946-4.

[5] T. Takaya, Y. Nihashi, T. Ono, H. Kagami, Transcription of endogenous retrovirus group K members and their neighboring genes in chicken skeletal muscle myoblasts, J. Poult. Sci. 58 (2021) 79–87. https://doi.org/10.2141/jpsa.0200021.

[6] S.S.M. Beski, R.A. Swick, P.A. Iji, Specialized protein products in broiler chicken nutrition: A review, Anim. Nutr. 1 (2015) 47–53. https://doi.org/10.1016/j.aninu.2015.05.005.

[7] W. Park, D. Rengaraj, D.Y. Kil, H. Kim, H.K. Lee, K. D. Song, RNA-seq analysis of the kidneys of broiler chickens fed diets containing different concentrations of calcium, Sci. Rep. 7 (2017) 11740. https://doi.org/10.1038/s41598-017-11379-7.

[8] M. Ghosh, N. Sharma, M. Gera, N. Kim, D. Huynh, J. Zhang, T. Min, S.S. Sodhi, M.B. Kim, V.P.B. Rekha, S. Ko, D. K. Jeong, Insights into phytase-containing transgenic Lemna minor (L.) as a novel feed additive, Transgenic Res. 27 (2018) 211–224. https://doi.org/10.1007/s11248-018-0068-z.

[9] S. Shinji, K. Umezawa, Y. Nihashi, S. Nakamura, T. Shimosato, T. Takaya, Identification of the myogenetic oligodeoxynucleotides (myoDNs) that promote differentiation of skeletal muscle myoblasts by targeting nucleolin, Front. Cell Dev. Biol. 8 (2020) 616706. https://doi.org/10.3389/fcell.2020.616706.

[10] J. Pohar, D. Lainscek, R. Fukui, C. Yamamoto, K. Miyake, R. Jerala, M. Bencina, Species-specific minimal sequence motif for oligodeoxyribonucleotides activating mouse TLR9, J. Immunol. 195 (2015) 4396–4405. https://doi.org/10.4049/jimmunol.1500600.

[11] A. Sanjaya, J.R. Elder, D.H. Shah, Identification of new CpG oligodeoxynucleotide motifs that induce expression of interleukin-1β and nitric oxide in avian macrophages, Vet. Immunol. Immunopathol. 192 (2017) 1–7. https://doi.org/10.1016/j.vetimm.2017.08.005.

[12] T. Takaya, Y. Nihashi, S. Kojima, T. Ono, H. Kagami, Autonomous xenogenic cell fusion of murine and chick skeletal muscle myoblasts, Anim. Sci. J. 88 (2017) 1880–1885. https://doi.org/10.1111/asj.12884.

[13] Y. Nihashi, T. Ono, H. Kagami, T. Takaya, Toll-like receptor ligand-dependent inflammatory responses in chick skeletal muscle myoblasts, Dev. Comp. Immunol. 91 (2019) 115–122. https://doi.org/10.1016/j.dci.2018.10.013.

[14] S. Ruijtenberg, S. den Heuvel S, Coordinating cell proliferation and differentiation: Antagonism between cell cycle regulators and cell type-specific gene expression, Cell Cycle 15 (2016) 196–212. https://doi.org/10.1080/15384101.2015.1120925.

[15] M.R. Daughtry, E. Berio, Z. Shen, E.J.R. Suess, N. Shah, A.E. Geiger, E.R. Berguson, R.A. Dalloul, M.E. Persia, H. Shi, D.E. Gerrard, Satellite cell-mediated breast muscle regeneration decreases with broiler size, Poult. Sci. 96 (2017) 3457–3464. https://doi.org/10.3382/ps/pex068.

[16] M. Hosotani, T. Kawasaki, Y. Hasegawa, Y. Wakasa, M. Hoshino, N. Takahashi, H. Ueda, T. Takaya, T. Iwasaki, T. Watanabe, Physiological and pathological mitochondrial clearance is related to pectoralis major muscle pathogenesis in broilers with wooden breast syndrome, Front. Physiol. 11 (2020) 579. https://doi.org/10.3389/fphys.2020.00579.

[17] K.J. Meloche, III W.A. Dozier, T.D. Brandebourg, J.D. Starkey, Skeletal muscle growth characteristics and myogenic stem cell activity in broiler chickens affected by wooden breast, Poult. Sci. 97 (2018) 4401–4414. https://doi.org/10.3382/ps/pey287.

[18] F.C. Meng, Z.F. Wu, Z.Q. Yin, L.G. Lin, R. Wang, Q. W. Zhang, Coptidis rhizoma and its main bioactive components: recent advances in chemical investigation, quality evaluation and pharmacological activity, Chin. Med. 13 (2018) 13. https://doi.org/10.1186/s13020-018-0171-3.

[19] S. Shinji, S. Nakamura, Y. Nihashi, K. Umezawa, T. Takaya, Berberine and palmatine inhibit the growth of human rhabdomyosarcoma cells, Biosci. Biotechnol. Biochem. 84 (2020) 63–75. https://doi.org/10.1080/09168451.2019.1659714.

[20] M. Imenshahidi, H. Hosseinzadeh, Berberine and barberry (Berberis vulgaris): A clinical review, Phytother. Res. 33 (2019) 504–523. https://doi.org/10.1002/ptr.6252.

[21] C. Bazzicalupi, M. Ferraroni, A.R. Bilia, F. Scheggi, P. Gratteri, The crystal structure of human telomeric DNA complexed with berberine: an interesting case of stacked ligand to G-tetrad ratio higher than 1:1, Nucleic Acids Res. 41 (2012) 632–638. https://doi.org/10.1093/nar/gks1001.

[22] M. Takagi, M.J. Absalon, K.G. McLure, M.B. Kastan, Regulation of p53 translation and induction after DNA damage by ribosomal protein L26 and nucleolin, Cell 123 (2005) 49–63. https://doi.org/10.1016/j.cell.2005.07.034.

[23] W. Jia, Z. Yao, J. Zhao, Q. Guan, L. Gao, New perspectives of physiological and pathological functions of nucleolin (NCL), Life Sci. 186 (2017) 1–10. https://doi.org/10.1016/j.lfs.2017.07.025.

[24] S. Soddu, G. Blandino, R. Scardigli, S. Coen, A. Marchetti, M.G. Rizzo, G. Bossi, L. Cimino, M. Crescenzi, A. Sacchi, Interference with p53 protein inhibits hematopoietic and muscle differentiation, J. Cell Biol. 134 (1996) 193–204. https://doi.org/10.1083/jcb.134.1.193.

[25] M.A. Cerone, A. Marchetti, G. Bossi, G. Blandino, A. Sacchi, S. Soddu, p53 is involved in the differentiation but not in the differentiation-associated apoptosis of myoblasts, Cell Death Differ. 7 (2000) 506–508. https://doi.org/10.1038/sj.cdd.4400676.

[26] A. Porrello, M.A. Cerone, S. Coen, A. Gurther, G. Fontemaggi, L. Cimino, G. Piaggio, A. Sacchi, S. Soddu, p53 regulates myogenesis by triggering the differentiation activity of pRb, J. Cell Biol. 151 (2000) 1295–1304. https://doi.org/10.1083/jcb.151.6.1295.

[27] S. Bauer, C.J. Kirschning, H. Hacker, V. Redecke, S. Hausmann, S. Akira, H. Wagner, G.B. Lipford, Human TLR9 confers responsiveness to bacterial DNA via species-specific CpG motif recognition, Proc. Natl. Acad. Sci. USA 98 (2001) 9237–9242. https://doi.org/10.1073/pnas.161293498.

[28] P.J. Bates, D.A. Laber, D.M. Miller, S.D. Thomas, J.O. Trent, Discovery and development of the G-rich oligonucleotide AS1411 as a novel treatment for cancer, Exp. Mol. Pathol. 86 (2009) 151–164. https://doi.org/10.1016/j.yexmp.2009.01.004.

[29] A. Slawinska, J. Brzezinska, M. Siwek, G. Elminowska-Wenda, Expression of myogenic genes in chickens stimulated in ovo with light and temperature, Reprod. Biol. 13 (2013) 161–165. https://doi.org/10.1016/j.repbio.2013.04.003.

[30] W. Luo, E. Li, Q. Nie, X. Zhang, Myomaker, regulated by MYOD, MYOG and miR-140-3p, promotes chicken myoblast fusion, Int. J. Mol. Sci. 16 (2015) 26286–26201. https://doi.org/10.3390/ijms161125946.

[31] H. Yue, X.W. Lei, F.L. Yang, M.Y. Li, C. Tang, Reference gene selection for normalization of PCR analysis in chicken embryo fibroblast infected with H5N1 AIV, Virol. Sin. 25 (2010) 425–431. https://doi.org/10.1007/s12250-010-3114-4.

